# Going back to ‘basics’: Harlow’s learning set task with wolves and dogs

**DOI:** 10.1101/2023.03.20.533465

**Authors:** Dániel Rivas-Blanco, Tiago Monteiro, Zsófia Virányi, Friederike Range

## Abstract

To survive and reproduce, animals need to behave adaptively by adjusting their behavior to their environment, with learning facilitating some of these processes. Despite the fact that dogs were the subject species for Pavlov’s original studies on learning, relatively little research has been done exploring dogs’ basic learning capabilities, and even fewer focused on the impact evolution may have had on this behavior. In order to investigate the effects of dog domestication on instrumental learning, we tested similarly-raised wolves and dogs in Harlow’s “learning set” task. In Experiment 1, several pairs of objects were presented to the animals, one of which was baited while the other was not. Both species’ performance gradually improved with each new set of objects, showing that they “learnt to learn” but no differences were found between the species in their learning speed. In Experiment 2 addressing reversal learning, once subjects had learned the association between one of the objects and the food reward, the contingencies were reversed and the previously unrewarded object of the same pair was now rewarded. Dogs’ performance in this task proved to be better than wolves’, albeit only when considering just the first session of each reversal, suggesting that either the dogs had not learned the previous association as well as the wolves or that dogs are more flexible than wolves. Further research (possibly with the aid of refined methods such as touchscreens) would help ascertain whether these differences between wolves and dogs are persistent across different learning tasks.

## Introduction

Learning is the process by which an animal acquires new knowledge, behaviors or skills (Gross, 2012) and proves to be crucial in the way animals interact with their environment: with past experiences informing current behavior, as well as information from novel opportunities (e.g., a new resource on which to feed) or risks (e.g., a poisonous animal which should be avoided) being integrated in their decision-making process and assisting survival.

One of the most common forms of learning is instrumental learning (also known as operant conditioning), defined as the association between a behavior and the outcome that results from it, often through a trial-and-error process (e.g., making an association between opening a garbage bin and finding food inside, (Bitterman, 1969). Instrumental learning is a highly conserved skill throughout taxa, being present in some capacity in animals ranging from nematodes to vertebrates (Gourgou et al., 2021; Pavlov, 1960). As such, it is particularly useful to design experimental paradigms that can easily be adapted across species with different cognitive abilities and *Umwelten* in order to compare their abilities (Uexküll & Mackinnon, 1926).

A classic paradigm used to test instrumental learning is the “learning set” paradigm - described by Harlow in 1949. In that study, rhesus macaques were initially tested in a discrimination task in which 344 pairs of different objects were sequentially presented. Of these, one object of each pair was considered the “positive” stimulus (which would grant a food reward upon being chosen by the subject), and the other, the “negative” one (which would beget no reward). Each object pair was presented several times so that the subject could learn the association between choosing the positive stimulus and getting the reward (Harlow, 1949). The subjects showed a gradual improvement throughout object pairs, getting an almost 100% success rate at trial 2 by the end of the experiment (i.e., after experiencing hundreds of object pairs). This showed that macaques were not only able to associate choosing the positive stimulus with getting the food reward, but to transfer the knowledge of the contingencies of the test to new iterations of it, therefore “learning to learn” (Shettleworth, 2010). After being tested in this initial task, the same subjects participated in a subsequent experiment in which the reward contingencies within each object pair were reversed (i.e., the previously negative stimulus would be the positive one and vice versa). As was the case with the preceding experiment, an almost 100% success rate at trial 2 was reached for the last reversals. Interestingly, even though this second experiment was, in principle, a more complex task, the subjects showed consistently better performance than in the previous experiment, possibly as a result of them generalizing from the discrimination task to the reversal (i.e., they not only “learned to learn” within the same task, but also between two similar tasks).

Similar experiments were later performed on other species (e.g., chimpanzees (Hayes et al., 1953), marmosets (Miles & Meyer, 1956), rats (Koronakos & Arnold, 1957), cats (Warren, 1966), pigeons (Zeigler, 1961), blue jays and crows (Hunter, 1970). However, the subjects rarely achieved that nigh-perfect level of success the rhesus macaques reached at trial 2 of each new object pair. In many cases, this could be attributed to the reduced number of object pairs the animals were trained to discriminate. Koronako’s and Arnold’s study (1957) shows that only a small fraction of subjects (5 of the 20 rats tested) were able to achieve 80% of correct choices for all 8 sets presented, although the compounding effect of each successive discrimination was not taken into consideration (i.e., whether they learnt to learn was not tested for, as no analysis were performed on their improvement with each successive set). Similarly, pigeons (Zeigler, 1961) reached around 60% of successful choices at second trial and around 70% on the last trial of the last few sets (80% when taking only the data from the most successful subjects), and although pigeons were exposed to many more sets of items than the rats, set numbers were still considerably less than in Harlow’s original study (with 120 sets presented versus 344 used in Harlow’s original experiment). After this original outburst of learning set experiments, later experiments focused on the “reversal” learning part of the paradigm, more as a measure of flexibility rather than of learning per se, often without being previously presented with a preceding sequential discrimination task, different to the original experiment (Bond et al., 2007; Erdsack et al., 2022; Rayburn-Reeves et al., 2013).

One species that is notoriously absent from the instrumental learning literature is dogs, with only a few studies done on reversal learning (Brucks et al., 2019; Frank, 2011; Frank & Frank, 1987; Tapp et al., 2003). This is particularly puzzling considering that not only were dogs the subjects for Pavlov’s original learning studies (Shettleworth, 2010), but also a widely studied species in the realm of animal cognition (particularly in the study of social cognition (e.g. Fugazza et al., 2016; Horowitz, 2009; Nagasawa et al., 2011), for reviews see (Bensky et al., 2013; Miklosi, 2007; Range & Marshall-Pescini, 2022). Given the interest in dogs’ cognitive abilities, the absence of studies on their learning skills is concerning, considering that it may be partially (or fully) responsible also for the performance observed in these experiments (Dickinson, 2012).

Compared to studies on dogs’ social cognition, few studies have addressed how domestication may have affected their learning skills. A study done by Frank and Frank (1987) in which both a discrimination and a reversal task were carried out showed that dogs outperform wolves when it came to reversal learning (but not to the basic discrimination learning). However, it has been pointed out that these wolves’ performance may have been an artifact of them being uncomfortable with the testing setting, as they were socialized to humans to a limited degree. Indeed, later on, this study was replicated with hand-reared wolves, which outperformed not only their mother-reared counterparts, but also dogs; both in the reversal task but on the discrimination one as well (Frank, 2011). A reversal learning study was also carried out in dogs and wolves by Brucks et al. (2019), but they did not find any differences between the species. It remains unclear, then, whether there truly are differences between these species’ learning skills.

One of the most relevant hypotheses has been put forward by Frank (2011) suggesting that domestication has largely increased the tractability of dogs by endowing them with a sensitivity to a broader band of stimuli (in particular, to arbitrary cues with no functional connection with the outcome) and with sufficient behavioral plasticity, preparing them to fulfill different jobs (e.g., police and sheep herding dogs). If so, one would expect dogs to be faster in Harlow’s instrumental learning tasks, in comparison with wolves (their closest-living relatives (Ostrander et al., 2019), who did not undergo the domestication process. Importantly, this is likely to include better performance in both phases of the task: higher responsiveness to arbitrary stimuli likely facilitates object-food associations whereas higher behavioral plasticity likely enables more flexible reversals. Beyond Frank’s tractability hypothesis, other findings have shown that wolves are, in general, more motivated and persistent than dogs (Marshall-Pescini, Virányi, et al., 2017; Rao et al., 2018), which gives them an advantage in problem solving tasks (Chow et al., 2016). Accordingly, it would be expected that wolves would outperform dogs in experiments testing their learning abilities by virtue of them being more engaged in the task (persistence hypothesis). This higher persistence, however, could prove detrimental in reversal learning tasks, as it could hamper their ability to switch strategies when the reward contingencies get swapped.

Furthermore, the “social-ecology” hypothesis (Marshall-Pescini, Cafazzo, et al., 2017) postulates that wolves’ and dogs’ different behaviors come as a result of adaptations to their socio-ecological environment. Wolves are cooperative hunters living in relatively stable (but also relatively scarce) environments, whereas dogs live in the comparably complex and ever changing human dominated niche. Over the process of domestication, dogs associated first with hunter gatherers, then with humans living in small settlements and finally with humans living in larger villages and cities. These ancient dogs (and, similarly, modern-day free-ranging dogs), may have access to plentiful food sources. However, these resources may nonetheless vary considerably across time and space (e.g., human refuse, garbage bins, fecal matter, etc., Atickem et al., 2009; Hughes & Macdonald, 2013; Lord et al., 2013; Mech et al., 2015; Mech & Boitani, 2007; Vanak & Gompper, 2009). Moreover, humans’ attitudes and behaviors towards dogs vary depending on various factors (e.g. state of affairs, age, local politics). As such, it would follow that dogs’ reliance on the changing human-shaped environment should make dogs more capable of discarding associations that are no longer adaptive, making them more flexible, even if not necessarily faster, learners than wolves.

This study aims to explore dogs’ and wolves’ performance on a task inspired by Harlow’s (1949) “learning set” task. Similar to Harlow’s study, we used several object pairs for the first discrimination experiment (i.e., the one in which one object was associated with a food reward) and, after a learning criterion was met, the subjects faced another experiment in which the same pair of items switched back and forth between baiting one item or the other (similar to Harlow’s reversal experiment). Critically, and to ease cross-species comparisons, dogs and wolves that participated in this study were raised and kept under similar circumstances.

## Methods

### Subjects

Experiment 1 started with 17 wolves (15.2 ± 2.4 months at first session) and 22 dogs (14.2 ± 2.4 months at first session). A subset of these animals was also tested in Experiment 2 (8 wolves (28.4 ± 3.5 months at the first session —see **Supplementary Table 1**) and 7 dogs (36 ± 4.6 months at the first session of acquisition).

Both wolves and dogs (**Figure 1a)** were raised in a similar environment. They were separated from their mothers at 10 days of age and then hand-raised by humans for 5 months. The pups were then integrated in packs with other adult conspecifics and housed in large 2000-8000m^2^ outdoor enclosures. All animals were trained to perform basic commands, participated regularly in behavioral experiments in the same testing facility, and had daily interactions with the experimenters. A complete list of subject related information can be found in **Supplementary Table 1**.

**Figure 1.**
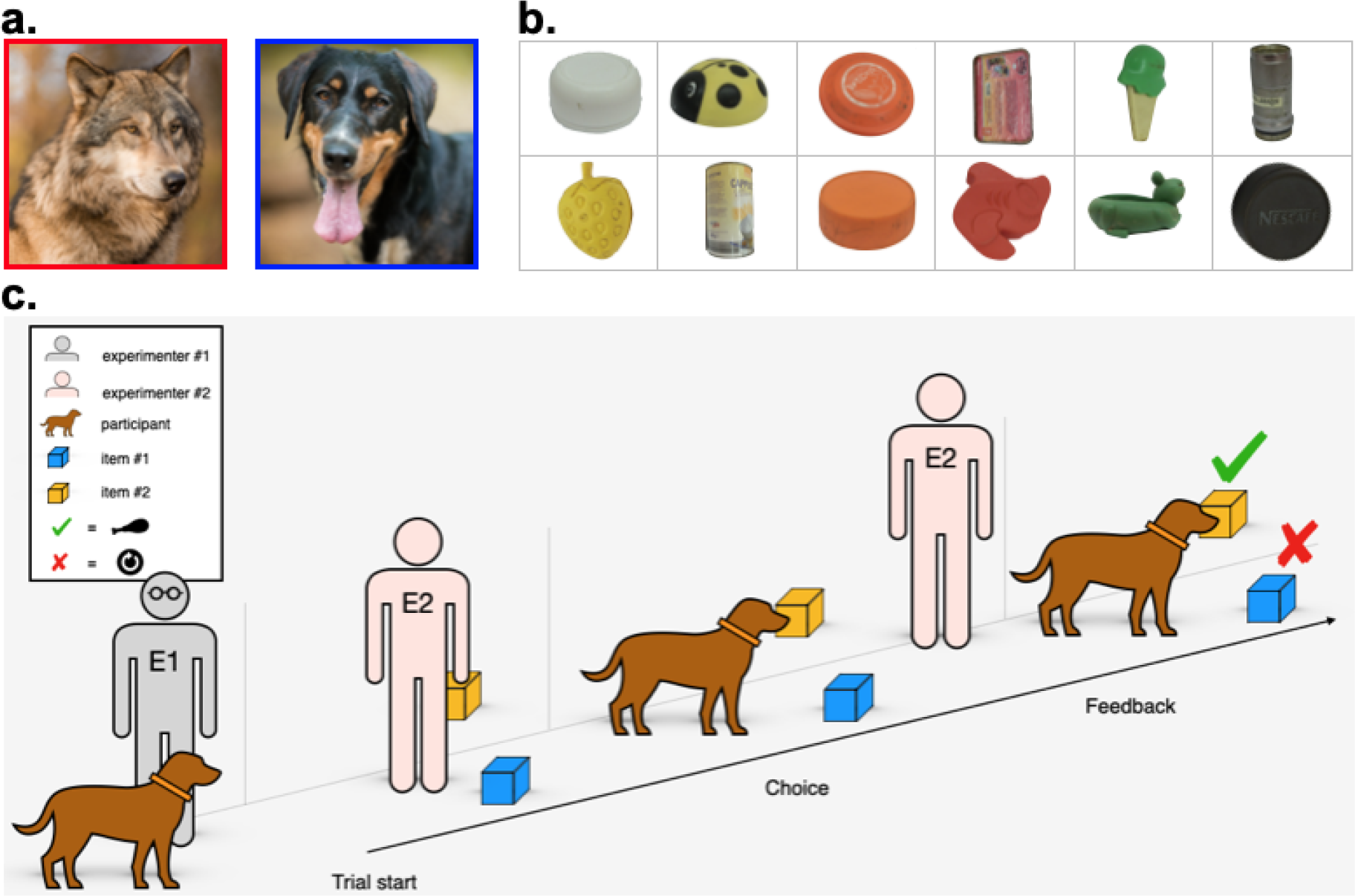
a. Subjects: Wolves (*Canis lupus*) and dogs (*Canis familiaris*) raised and kept in similar environments participated in this study. See *Supplementary Table 1* for details. **b**. Stimuli: Example items used in the experiments. See *Supplementary Table 2* for details **c**. Procedure: two human experimenters (E1 and E2) were needed to run Experiments 1 and 2. While E1 controlled the participants, E2 was responsible for stimuli placement and associated contingencies. The trials of both experiments consisted of 3 main parts: trial start, choice, and feedback. See *General procedure* for details.

### Materials

Several pairs of items of different sizes and colors were presented to the subjects, all of them with some sort of crevice or gap under which a piece of high-quality reward (meat or sausage) could be placed. A comprehensive list of the items used can be found in **Supplementary Table 2**, example pictures can be found in **Figure 1b**.

### Testing facility

Testing was conducted in a large indoor testing area of 5.4 × 9 m. To reduce the possibility of the animals developing side bias, the starting position of the animals was changed on every trial within a session, and the objects were placed at different locations (i.e., moving to different areas within the testing chamber).

### General procedure

Both Experiments 1 and 2 shared the same basic procedure (**Figure 1c**). Two experimenters (E1 and E2 hereafter) conducted the study. E1 called the animal to the starting position, asked it to sit down next to them, rewarded it with a food reward and then held the animal by the collar. E2 baited the positive stimulus with a food reward (baiting was done with their back turned so that animals could not see it), and thoroughly smeared the second stimulus with food to prevent the animals from finding the food reward by their sense of smell alone. Then E2 moved to stand in front of E1 with their back facing the subject, holding both objects in their hands. The distance between E2 and the subject at the start of each trial would depend on the species, 1 long step (∼1m) for dogs and 2-3 steps for wolves due to the differences in size and speed between the species (except for one of the wolves —Nanuk—, for which the distance was reduced to 1-1.5m due to his sight impairment).

Once E2 was in position, E1 either stared down or closed her eyes to avoid cuing the animal inadvertently. E2 then stepped about 2-3 meters forward and placed the two objects on the floor one after the other, around 1.5-2m away from each other, by extending their arms and crouching, but keeping their body approximately at the same distance from both objects. After this, E2 took another step forward, leaving the objects behind them, with their back still facing the animal.

E1 then released the animal and the trial commenced. A subject would be considered to have made a choice when they came into contact with one of the objects. If the subject made the correct choice, they were allowed to eat the reward. However, if the subject made the incorrect choice, E1 would walk towards the reward under the positive stimulus and take it away before the animal could reach it (due to safety concerns, in cases in which E1 would not be able to take away the reward, E2 would do it instead). If the subject failed to make a choice, the trial was repeated until they did so. However, some subjects lost motivation during some sessions and failed to make a choice, regardless of the repetitions of that trial. In such cases, the session was aborted on that trial.

Each session consisted of 10 trials, in 5 of them the positive item was on the right side, in 5 of them on the left side; no more than twice in a row on the same side. The order in which the items were placed on the floor was counterbalanced according to a predetermined assignment, with the positive stimulus presented first on some trials and second on others; this ordering was also not repeated more than twice in a row within the same session. Furthermore, the same combination of side and order of presentation was never repeated (i.e., if the positive stimulus was presented first on the right side in a given trial, the next trial in which the positive stimulus was placed on the right side, it would have to be presented second).

Sessions were carried out once a week, but larger gaps took place whenever other experiments were carried out (as well as any other disruptive events, such as health checks and the breeding season).

### Experiment 1: Discrimination learning

During this Experiment, several sets of items were presented to the subjects (**Supplementary Table 2**). Within sets, items were of different colors. Furthermore, color pairings used in a given set of items could not be repeated within the subsequent 5 sets. Whenever a subject was exposed to an item with a color that matched another item from a previous set, this item was of the opposite stimulus type (i.e., if a purple item was used as a positive stimulus in one of the sets, the next time a purple item was used for the same subject, it was the negative stimulus instead). This counterbalancing was done to prevent spontaneous biases from potentially arising within the subjects, as dogs have been shown to be able to discriminate between a variety of colors despite only having two different types of cone photoreceptor cells (Kasparson et al., 2013). Nonetheless, some of the items used were considered as “mixed” color (i.e., they did not have a unique color taking most of their surface area, see **Figure 1b** and **Supplementary Table 2**), and the above-mentioned restrictions did not apply to them.

Each time a subject chose the positive stimulus in 9 out of 10 trials within a session, this session was considered successful and a new set of items was presented in the subsequent session. If out of 5 sets in a row 4 were successfully completed in the first session and 1 set was completed within the first two sessions, the subject was considered to have reached the learning criterion and moved on to Experiment 2 (see below).

### Experiment 2: Reversal learning

The procedure was similar to Experiment 1, but only a single set of items was presented to the subjects. After an initial successful “acquisition” session (9 out of 10 correct trials), the reward contingencies associated with each stimulus would be switched (i.e., the original positive stimulus becoming negative, and vice versa). Stimulus reward contingencies would be reversed after each successful session (9 out of 10 correct trials; except for three occurrences in which a subject moved on to the next reversal with less correct trials due to human error —Aragorn’s 10^th^ reversal and Amarok’s 8^th^ reversal, with 8 out of 10 correct trials; and Nuru’s 3^rd^ reversal with 7 out of 10). These reversals would take place until the learning criterion was achieved (i.e., out of 5 consecutive reversals, 4 of them needed to be completed within the first session, and 1 within either the first or second session).

### Data selection

The first 15 sets (Experiment 1) were used for data analysis. Since different subjects reached criterion at different speeds (and some did not reach it), we decided to consider only the first 15 sets, a point before which none of the subjects had moved to *Experiment 2* (see **Supplementary Table 1** for more information on the performance of each subject). To make the size of the dataset similar to that of Experiment 1, we used only the first 15 reversals for Experiment 2. We considered this our initial dataset, and all of the additional data selection criteria described below were performed upon this dataset.

Additionally, we removed all sessions in which experimental errors or anomalies took place from the analyses (4.8% of the dataset). Examples of these situations include sessions done after the 9/10 criterion was met (due to human error), sets that were not completed (because the items got broken), and sessions with more than two null trials (because the animals were not motivated to participate in the test). Whenever a session or a set was removed according to this criterion, the number of the session or set was transferred to the next valid one (e.g., if an animal’s 4^th^ set was removed from the dataset, the following set would be considered set 4).

Furthermore, for a subset of the subjects in a reduced number of sessions (0.9% of the dataset) the procedure was considerably different (i.e., the subject stayed in a room adjacent to the testing chamber while E2 placed the items and would later be released back into the main testing chamber through a sliding door). These sessions were excluded from analyses as well.

Sessions that did not take place between March of 2009 (the beginning of the experiment) and February of 2016 were also excluded from the analyses (3.4% of the dataset). After February of 2016, sessions would take place once every few weeks for each subject, but afterwards larger gaps of several months would be commonplace for all individuals. As increased and variable gaps in the testing regime could affect performance in a way not accounted for by our experimental design, we decided to exclude these sessions as well.

All sessions that were removed from analyses are still included in the dataset presented (**dataset 1**; see also **Supplementary Table 1** for more information on the performance of each subject), alongside with the criterion that warranted their exclusion.

### Data analysis

We used R version 4.0.4 (R Core Team, 2021); https://www.r-project.org) to carry out all statistical analyses. For each experiment, two binomial Generalized Linear Mixed Models (GLMM) with logit link function were conducted to analyze the data (lme4 package; Bates et al., 2015).

The response variable for the first group of models (**Table 1**, models 1 and 3) was the outcome of the session (i.e., whether the criterion of 9 out of 10 correct responses was met). The main predictors were the interaction between species and session (i.e., whether the species improved at a different speed within the sessions of the same set) and the interaction between species and set—in Experiment 1— or reversal number —in Experiment 2— (i.e., whether the species improved at a different speed along the sets and within the same session in those sets). To account for the fact that sessions and sets had a nested structure (with each session being part of a single set/reversal), we created an ID for each session within a given set/reversal and included this variable (nested within set/reversal number) as a random intercept of the model. Finally, we also included the subjects as a random intercept.

**Table 1.**
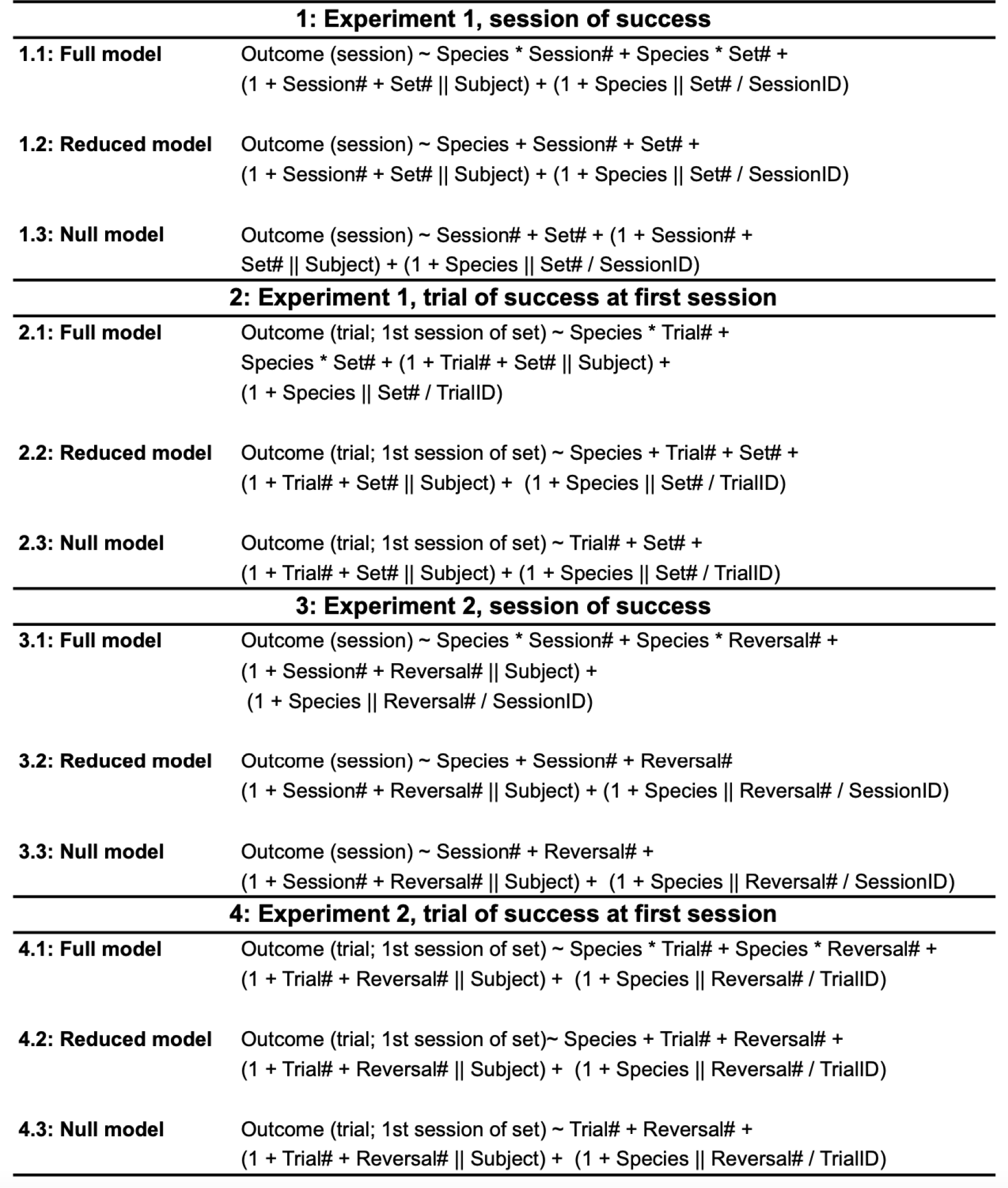
Statistical models.

Since these models did not distinguish between performances below the 9 out of 10 criterion (e.g., a session with 8 out of 10 correct trials would be considered no different from 2 out of 10 correct trials), and in order to account for the fact that different subjects would perform a different amount of trials within each set or reversal (as they needed more or less sessions to reach criterion within each set/reversal), we decided to fit a second group of models (**Table 1**, models 2 and 4). In these models, the response variable was the outcome of each trial (success or failure) within the first session of a given set/reversal, and the predictors were the interaction between species and trial (i.e., learning speed within each set for each species) and the interaction between species and set number (learning speed within each trial for each species). Similarly to the previous set of models, we created an ID for each trial within each set/reversal and included it as a random intercept nested within the number of the set/reversal. The subjects were also included as a random intercept.

All numerical variables were z-transformed for model fitting. Random slopes were added whenever the variable had at least three different values per level of the respective random intercept (for continuous variables) or at least two levels with at least two observations per level of the respective random intercept (for discrete variables).

Each of these models was compared with a *null* version that had the same structure but excluded species as a variable. To account for the possibility that there were no differences in performance as a factor of improvement along trials, sessions, or sets/reversals, but still overall differences in performance between the species (as well as for the possibility that there are no differences between the species, but still variation along trials, sessions, or sets/reversals), a *reduced* version of the models (with the same predictors as the *full* models, but without the interaction) was also fitted and subsequently compared with the null as well (**Table 1**).

All models were tested for collinearity with the *vif* function from the *car* package (Fox & Weisberg, 2018) and further tested for model stability through a set of custom-made functions, designed by Roger Mundry and later edited by Remco Fokertsma, and the *DHARMa* library (Hartig, 2020). All models met stability and colinearity assumptions, with variance inflation factors (vif) under 3 in all cases (largest vif = 2.931; Field, 2009; Quinn & Keough, 2002).

### Ethics statement

No special permission for use of animals (wolves and dogs) in socio-cognitive studies is required by Austrian law (Tierversuchsgesetz 2012–TVG 2012). The relevant committee that allows running research without special permissions regarding animals is: Tierversuchskommission am Bundesministerium für Wissenschaft und Forschung (Austria).

## Results

### Experiment 1: Discrimination learning

When looking at the animals that made it to at least set 15 (15 out of the original 22 dogs and 16 out of the original 17 wolves, see **Supplementary Table 1** for more details), both species needed a similar number of sessions to reach set 15, with 43.6 sessions on average for dogs, and 43.1 sessions on average for wolves.

With each new set of objects presented, the subjects became faster at associating the food reward with choosing the positive stimulus (needing less sessions per set to reach the criterion; **Figure 2**). However, we did not find differences between the species (full-null comparison —**Table 1:** 1.1 and 1.3—: χ^2^ = 0.456, p = 0.928; reduced-null comparison **Table1:** 1.2 and 1.3—: χ^2^ = 0.221, p = 0.638). As for the null model, (**Table 1:** 1.3), both the effects of the session number (estimate ± SE: 0.547 ± 0.105; z value = 5.215, p < 0.001) and the set number (estimate ± SE: 0.303 ± 0.059; z value = 5.097, p < 0.001) were significant, showing that, indeed, subjects needed less sessions to reach criterion in later sets, and that each session within the same set increased the likelihood of success.

**Figure 2.**
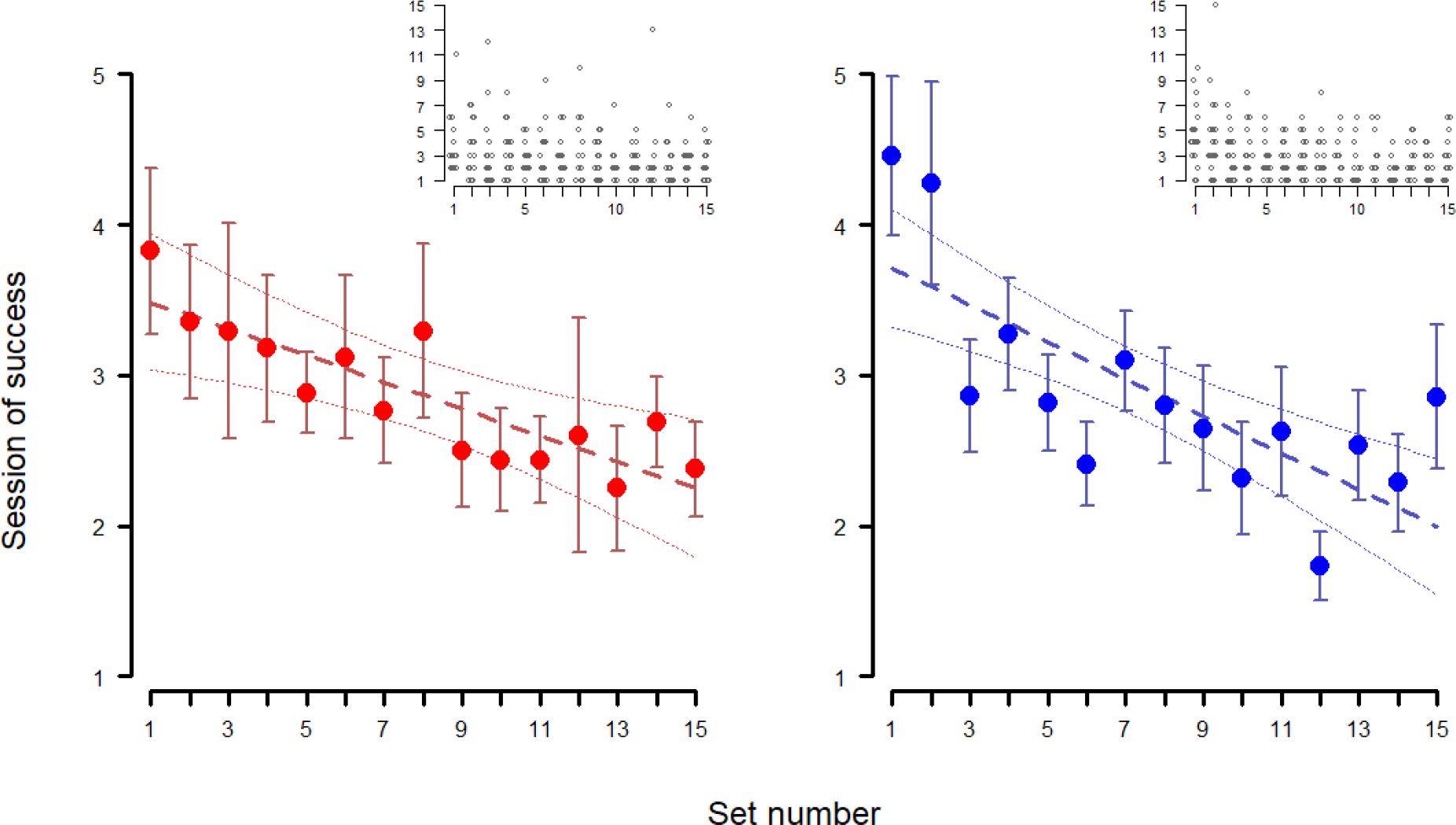
Across-session discrimination behavior - Experiment 1. Average session of success at each set (at least 9 out of 10 correct trials) in Experiment 1 for wolves (red, left) and dogs (blue, right). Whiskers represent the standard error. Regression lines were calculated through a linear model with session of success as the dependent variable and set number as the independent variable, with confidence intervals set at 95%. Insets show individual data points.

When analyzing the outcome of each trial on the first session of each set (**Table 1:** 2.1), once again, we did not find an effect of the species (full-null comparison —**Table 1:** 2.1 and 2.3—: χ^2^ = 0.547, p = 0.908; reduced-null comparison —**Table 1:** 2.2 and 2.3—: χ^2^ = 0.517, p = 0.772), as can be seen in **Figure 3**, for sets 1, 7 and 15. Similarly to the previous model, both the trial number (estimate ± SE: 0.206 ± 0.022; z value = 9.454, p < 0.001) and the set number (estimate ± SE: 0.085 ± 0.031; z value = 2.778, p = 0.005) were significant, meaning that, when looking at the first session per set, both species improved along trials and sets.

**Figure 3.**
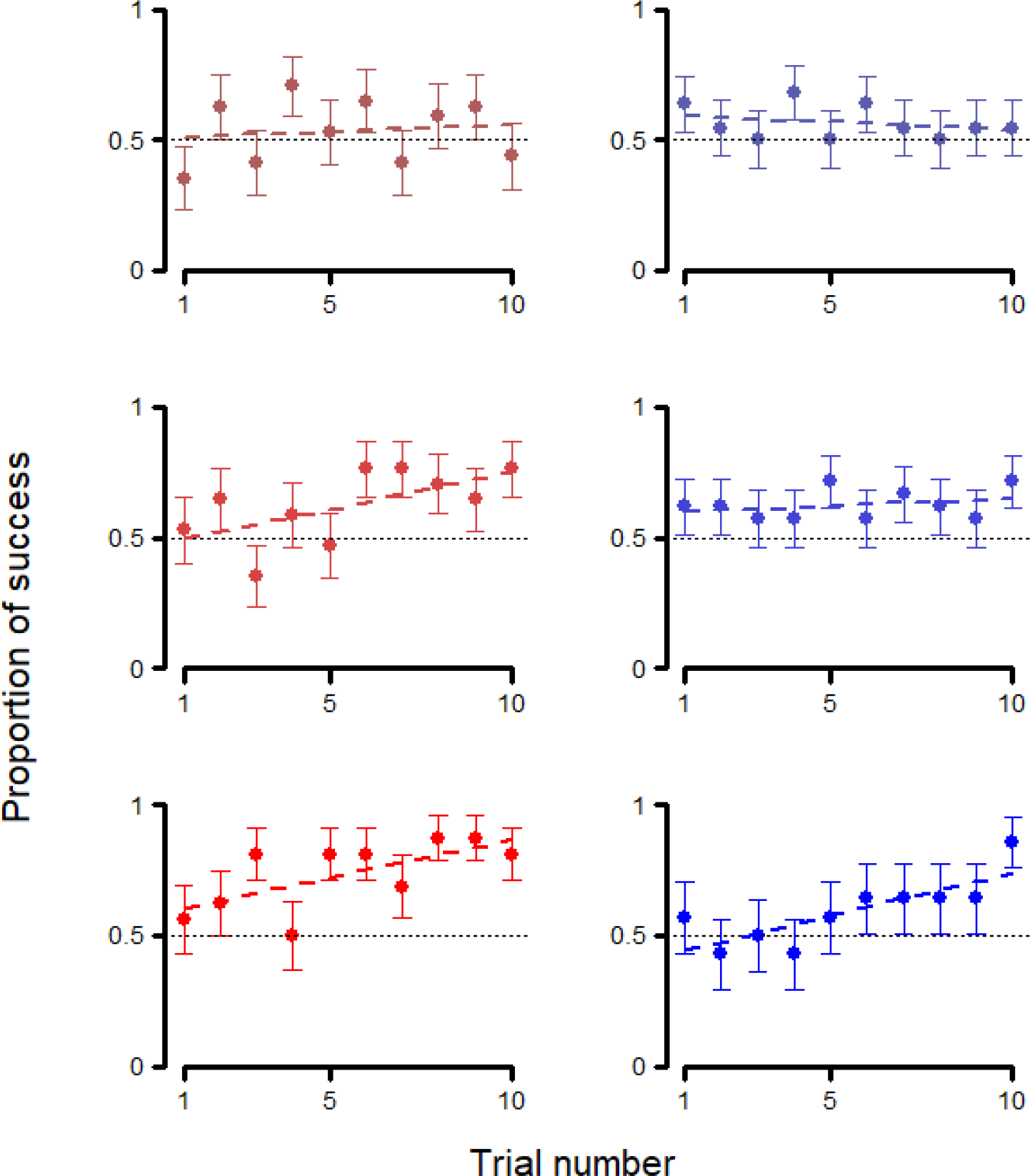
Trial-by-trial behavior at first session - Experiment 1. Average performance in each trial of Experiment 1 at the first session of sets 1 (1^st^ row), 7 (2^nd^ row), and 15 (3^rd^ row), for wolves (left, red) and dogs (right, blue). Whiskers represent the standard error. Regression lines were calculated through a linear model with session of success as the dependent variable and trial number as the independent variable.

### Experiment 2: Reversal learning

The subjects that continued on to Experiment 2 (6 dogs, 8 wolves; see **Supplementary Table 1** for more details) had experienced a similar number of sets in Experiment 1, with dogs going through an average of 32.8 sets and wolves through an average of 34.0 sets. They also had done a similar total number of sessions in Experiment 1 with 80.17 sessions for dogs and 78.3 sessions for wolves. Out of these subjects, 3 dogs and 7 wolves completed 15 reversals, with the average number of sessions to do so being 34.3 for the dogs and 46.0 for the wolves.

Similarly to Experiment 1, we did not find any differences between the species when analyzing the session at which they reached criterion each time the reward contingencies were reversed (full-null comparison —**Table 1:** 3.1 and 3.3—: χ^2^ = 2.503, p = 0.475; reduced-null comparison —**Table 1:** 3.1 and 3.3—: χ^2^ = 0.799, p=0.371). Furthermore, there was no apparent learning effect along successive reversals for either species (see **Figure 4**). This was also shown in the models, as there was no significant effect of the reversal number in the null model (**Table 1:** 3.3—: estimate ± SE: 0.041 ± 0.156; z value = 0.263, p = 0.793), although there was still an effect of the session number (estimate ± SE: 1.428 ± 0.170; z value = 8.359, p < 0.001). Taken together, these results indicate that the animals did not get significantly better with each reversal, or at least, that they did not improve enough with each reversal so as to reduce the amount of sessions needed to reach the 9 out of 10 success criterion.

**Figure 4.**
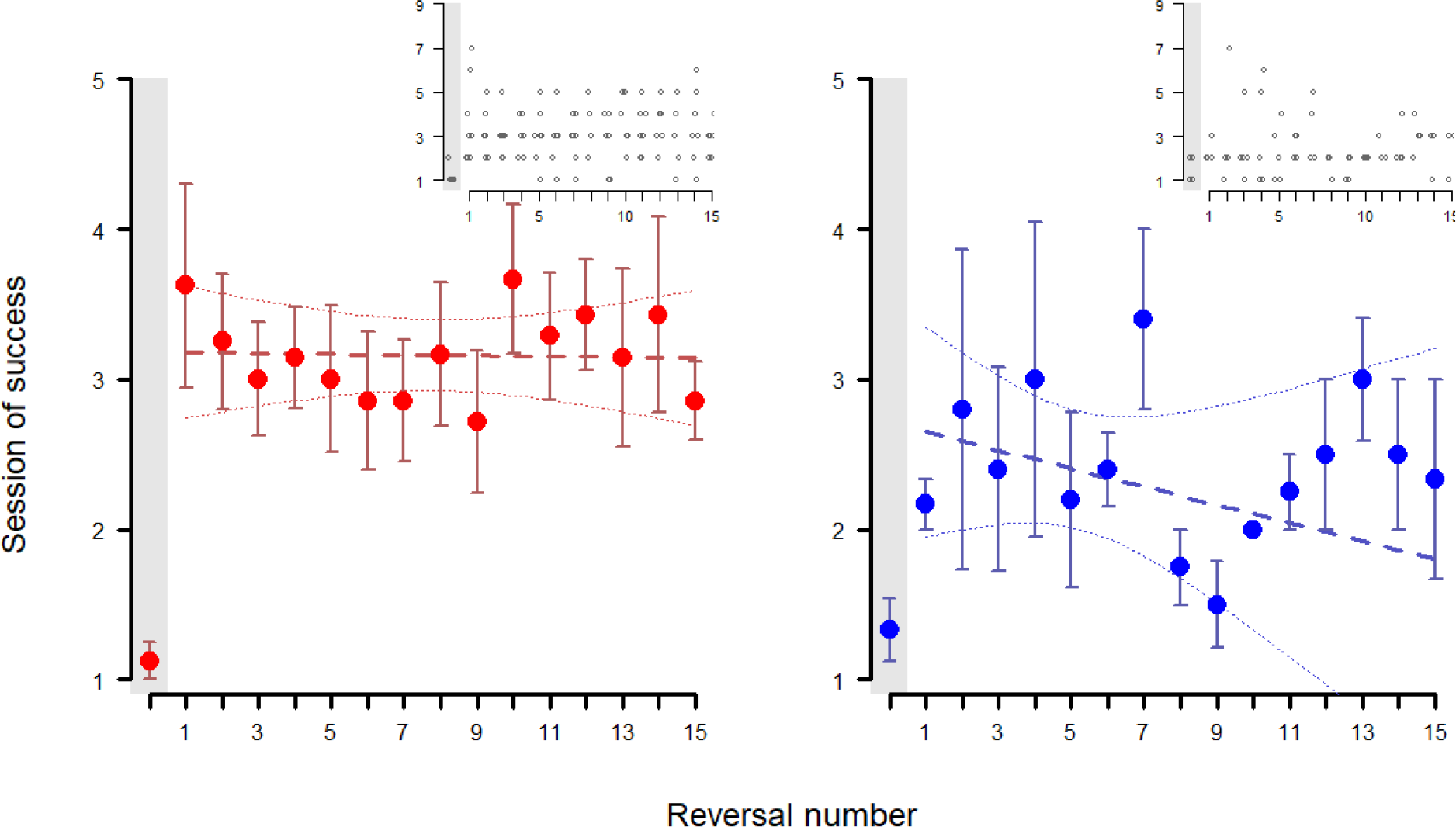
Across-sessions discrimination behavior - Experiment 2. Average session of success at each reversal (at least 9 out of 10 correct trials) in Experiment 2 for the wolves (left, red) and the dogs (right, blue). “Reversal 0” (highlighted in gray) refers to the acquisition of the association, before the reward contingencies were reversed (functionally equivalent to one of the sets in Experiment 1). Whiskers represent the standard error. Regression lines were calculated through a linear model with session of success as the dependent variable and reversal number as the independent variable, with confidence intervals set at 95%. Insets show the individual data points.

However, there was a significant effect of the species on the outcome of the trials on the first session of each reversal (full-null comparison —**Table 1:** 4.1 and 4.3—: χ^2^ = 12.977, p = 0.005), with the improvement with each passing trial being significantly worse in wolves (estimate ± SE: -0.158 ± 0.070; z value = -2.247, p = 0.025; **Figure 5**), although the interaction between species and set number was not significant (estimate ± SE: 0.099 ± 0.080; z value = 1.235, p = 0.217).

**Figure 5.**
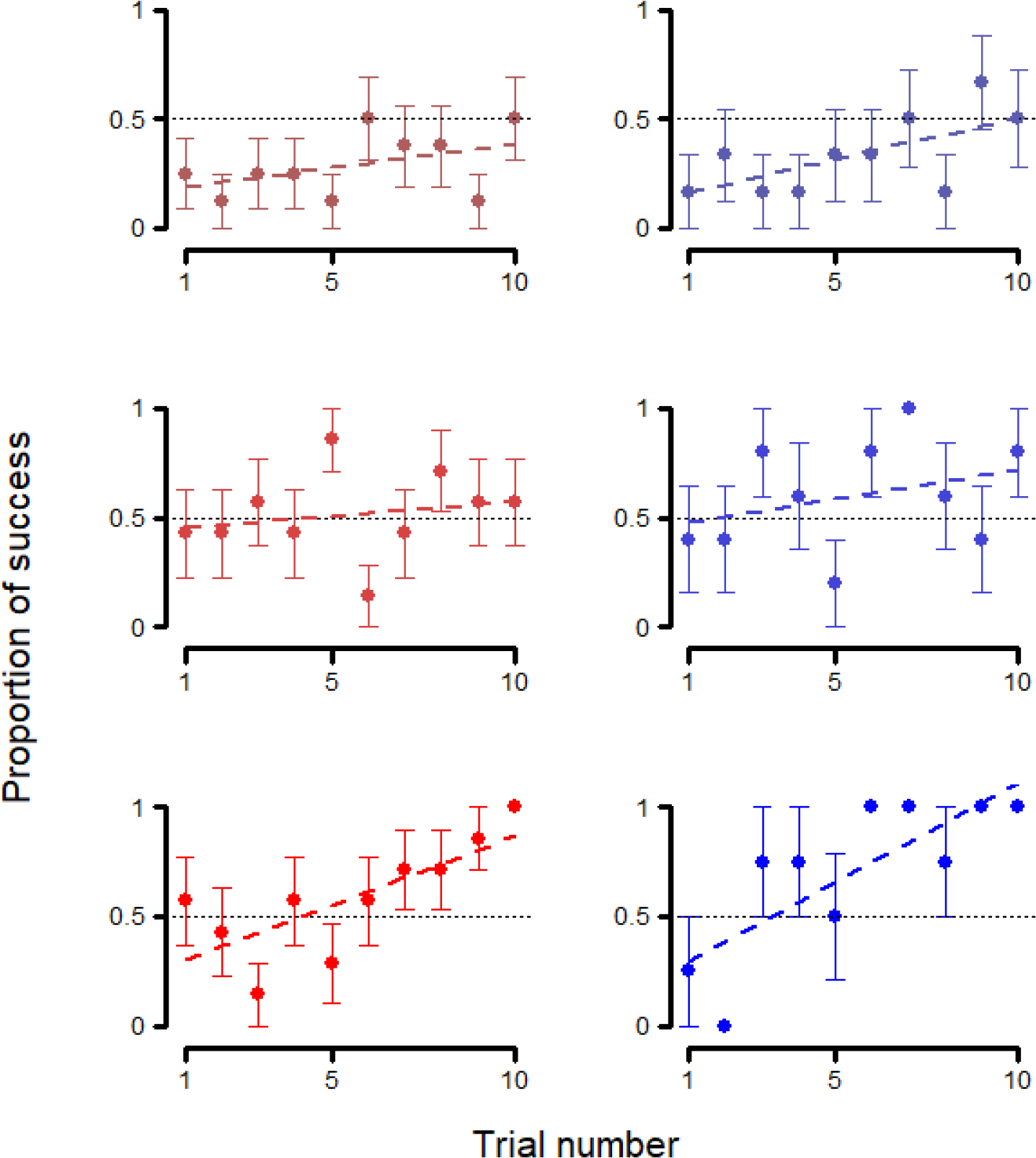
Trial-by-trial behavior at first session - Experiment 2. Average performance in each trial of Experiment 2 at the first session of reversals 1 (1^st^ row), 7 (2^nd^ row), and 15 (3^rd^ row), for wolves (left, red) and dogs (right, blue) Whiskers represent the standard error. Regression lines were calculated through a linear model with session of success as the dependent variable and trial number as the independent variable.

We still found an effect of species when comparing this reduced model with the null model (reduced-null comparison —**Table 1:** 4.2 and 4.3—: χ^2^ = 6.336, p = 0.012; estimate ± SE: -0.279 ± 0.101; z value = -2.769, p = 0.006), and in this reduced model, also the trial number (estimate ± SE: 0.400 ± 0.034; z value = 11.690, p < 0.001) and the reversal number (estimate ± SE: 0.191 ± 0.049; z value = 3.877, p < 0.001) were significant suggesting that the subjects’ performance on the first session did improve each time the reward contingencies were reversed.

## Discussion

In this study we examined wolves’ and dogs’ instrumental learning skills in both a serial discrimination and a reversal learning task. We did not find any differences between the species in the acquisition of associations (Experiment 1), but the performance of each species did improve with each additional set of stimuli presented, showing that they both have “learnt to learn”. However, dogs did perform better than wolves in Experiment 2 —where the task was to extinguish learned associations and instead acquire an association with the previously unrewarded stimulus. However, dogs only outperformed wolves when analyzing the first session of each reversal, but no differences were found between the species along reversals. Similarly, dogs did not reach session criterion within each reversal faster than wolves, and neither species showed improvement at the session level (i.e., they did not need fewer sessions to achieve criterion with each reversal).

Taken together, our results seem to support the “social ecology” hypothesis which predicts similar learning speed in dogs and wolves but higher flexibility in dogs than in wolves (Frank, 2011; Marshall-Pescini, Cafazzo, et al., 2017). As dogs often live in rapidly changing human-shaped environments, the ability to discard learned behavior that is no longer useful is particularly important for survival within their ecological niche (Marshall-Pescini, Cafazzo, et al., 2017). In line with this, in the current study only the second experiment showed any differences between the species (even if only to a limited degree). The finding that dogs were not consistently “faster learners” than wolves seems to contradict Frank’s “tractability” hypothesis —which postulates that, as dogs were selected to perform many different tasks for humans, they should be faster at making new associations as well as their corresponding reversals (Frank, 2011).

We cannot exclude, however, the possibility that dogs are, as predicted by the tractability hypothesis, better capable of making arbitrary associations for which wolves can, however, compensate with their higher persistence. Wolves have been shown to be more persistent than dogs, whereby they engage in problem solving tasks for longer than dogs and they are not deterred even when no solution is available (Rao et al., 2018). Thus, it could be the case that, in Experiment 1, wolves’ persistence and dogs’ affinity towards tasks involving arbitrary cues lead to their comparable success and learning speed. In Experiment 2, however, a higher level of persistence no longer supported success, as the reward contingencies were regularly switched, thus in this task the higher behavioral plasticity of dogs could become apparent.

Nevertheless, it remains unclear why the observed increase in performance in the first sessions of each reversal seen in Experiment 2 did not translate into differences in learning speed between the species overall. One explanation for the diverging results of two models used to analyze the data in Experiment 2 would be the steep success criterion. As only one mistake was allowed in order to proceed to the next reversal, better performances within each session would not necessarily mean achieving the criterion faster. Non-successful sessions did range from 0 to 8 out of 10 correct trials (as opposed to the 9 or 10 out of 10 correct trials needed for a successful session), which could mean that differences in learning speed could have been obscured by the success criterion. This may have compounded with the fact that the task was more difficult than the one in Experiment 1, which could explain why such an effect was not an issue there. Indeed, neither species showed a decrease in sessions needed to reach criterion within a reversal even though they did get better at solving the task within the first session of each successive reversal, which may point to a possible floor effect derived from the criterion we used.

Regardless of the reason behind the inconsistency between the first session and overall performance within Experiment 2, it is also unclear what cognitive processes may underlie the observed differences in reversal learning between the species. Both the social ecology and the tractability hypotheses predict higher behavioral plasticity in dogs than in wolves, even if due to different selection pressures (frequent, human-induced changes in environment vs. diverse tasks and direct interactions with humans, respectively). What kind of changes in cognition and behavioral regulation support dogs’ flexible behavior, remains an open question. In our task, in order to acquire a new and opposite association, the previous one would have to be extinguished, and in order to do so, the previous response towards the positive stimulus would need to be inhibited (Shettleworth, 2010). In light of our results, it would be plausible that either the dogs were better than the wolves at inhibiting the learned association, or that the dogs made weaker associations (i.e., they did not remember the reward contingencies as well as the wolves did), and thus had an easier time “un-learning” them. However, we cannot tease these possibilities apart within the confines of the current study, as we did not include any controlled measures of memory in our paradigm. Dogs have been shown to retain learned associations for a period of at least 6 months (Wallis et al., 2016), making it unlikely that the observed results may come as a consequence of poor memory. Additionally, no differences in inhibition have been found between dogs and wolves (or, when they have, they were not extrapolatable among tasks (Brucks et al., 2019; Wallis et al., 2016), making it hard to draw any conclusions about any differences in the capability to inhibit learned associations. Thus, future studies need to measure both instrumental learning and memory in a controlled manner (e.g., as part of a battery of tests) in order to provide insight about the mechanisms driving the observed differences in reversal learning.

When compared to Harlow’s original study with the Rhesus monkeys, we did not find a nigh-perfect performance at trial two in our subject species (Harlow, 1949). However, this is, in all likelihood, a result of the reduced number of sets used in our study, when compared with Harlow’s 344 sets. Our number of sets was more comparable with Zeigler’s (1961) experiment on pigeons and Arnold’s (1957) experiments on rats. which reported performances above chance, but nevertheless not almost perfect by the second trial of each set. Nonetheless, even though comparisons between Harlow’s and ours are complicated (not only because of the differences in methodology, but also the manner in which the data was reported in the original study), it could be argued all things equalled, the performance by set 15 is not qualitatively dissimilar between our study and Harlow’s. In Harlow’s study, the average performance in sets 9 to 16 described a mostly linear curve starting at 50% of correct choices and rising to somewhat above 80% by trial 6. In our study, performance in the first session also starts indistinguishable from chance in trial 1 and then rises to around 75%. That is, of course, not to say that we should necessarily expect similar performances in non-human primates and canids were we to increase the testing effort.

Learning remains underexplored in dogs and wolves, with our results still failing to provide a clearer picture when put into context with the current literature. This further highlights the relevance of carrying out more experiments on this topic, as it is now apparent that much is still unknown about such an important process, which could prove fundamental to other studies performed on these species. It is our view that, without a solid grasp of how instrumental learning works in these species, the efforts towards studying other, more complex, behaviors that may rely on it may prove to be misguided.

One possible avenue to study instrumental learning in wolves and dogs would be to employ methods such as touchscreens or other automatized apparatuses. Possibly because the subjects do not need to be repositioned after each trial and the fact that the reward is dispensed automatically, touchscreen-based paradigms have been fairly successful in carrying out large volumes of trials in dogs and wolves (e.g., Aust et al., 2008; Dale et al., 2019; Laude et al., 2016; Range et al., 2008; Rivas-Blanco et al., 2020). As all touchscreen tasks need an instrumental learning training phase in order to teach the animals how to interact with the apparatus (usually by presenting them with a positive stimulus that is rewarded and a negative stimulus that is not, see (Rivas-Blanco et al., 2020), it should be fairly straightforward to extend this training regime to further sets of images.

In conclusion, we found that both wolves and dogs improve with repeated exposure to the paradigm (although never at near-perfect levels by trial 2), but roughly at the same speed. Dogs may outperform wolves when it comes to extinguishing previously made associations and learning new ones in their stead, but more research needs to be done on the matter. Further studies are needed that consider the use of automatized testing methods, as they allow for a higher testing volume while reducing the subjects’ fatigue.

## Supporting information

Supplementary Table 2 (objects)

Supplementary Table 1 (subjects)

## Supplementary materials

-**Supplementary Table 1 (subjects)**

List of all the subjects that participated in this study, with their species and sex, whether they participated only in Experiment 1 or both, their age at the beginning of Experiment 1 (and Experiment 2, whenever it applied) and the amount of sets and reversals they were exposed to in each experiment. Subjects marked in gray fulfilled the criterion of either the Experiment 1 (whenever the grayed out area reached the third-to-last column) or both experiments (whenever the whole row is grayed out).

-**Supplementary Table 2 (objects)**

IDs of the objects used in this experiment, together with their colors.

### Dataset 1

Dataset used for data analysis and in figures.

## Acknowledgements

We thank the trainers and students at the Wolf Science Center for carrying out the experiments and collecting the data. We also thank Gwen Wirobski and Andreas Berghäenel for comments on earlier versions of the manuscript and all members of the Domestication Lab for feedback during the course of the project.

## Data accessibility

All data generated or analyzed during this study are included in this published article as supplementary materials.

## Author contributions

ZV and FR conceptualized and designed the experiments; DRB curated the dataset; DRB and TM processed and analyzed the data; DRB, TM and FR wrote the first draft of the paper; all authors edited and reviewed the manuscript.

