## Supplementary Table 2 (objects) for "Going back to ‘basics’: Harlow’s learning set task with wolves and dogs"

| **Item** | **Color** |
|---|---|
| pink_barbie_box | pink |
| pink_barbie_box_lid | pink |
| black_ashtray | black |
| black_cofee_pot | black |
| black_cup | black |
| black_devil | black |
| black_filter | black |
| black_wheel | black |
| blue_castle | blue |
| blue_filter | blue |
| blue_havaball | blue |
| blue_pretzel | blue |
| blue_seahorse | blue |
| blue_strawberry | blue |
| blue_train | blue |
| brown_cappuccino_box | brown |
| brown_leaf | brown |
| brown_pot | brown |
| brown_cardboard_box | brown |
| white_pot_lid | white |
| cookie_box_lid | mixed |
| yellow_pedros_lid | yellow |
| green_croissant | green |
| green_face | green |
| green_plane | green |
| green_shovel | green |
| green_soap_frog | green |
| green_whale | green |
| grey_egg_holder | grey |
| grey_rubber_dogbasket_foot | grey |
| ice_cream_pot | mixed |
| silver_cup | silver |
| orange_castle | orange |
| orange_face | orange |
| orange_ovomaltine_box | orange |
| purple_big_half_ball | purple |
| pink_bowl | pink |
| purple_half_ball | purple |
| purple_multivitamin_tube | purple |
| red_bread | red |

| | |
|---|---|
| red_frisbee | red |
| red_hand | red |
| red_toy_horse | red |
| red_pear | red |
| red_shark | red |
| red_train | red |
| santa | mixed |
| silver_shopping_cart_wheel | silver |
| silver_coffee_pad | silver |
| silver_dish | silver |
| silver_pot | silver |
| silver_tube | silver |
| stone | mixed |
| white_dish_with_hole | white |
| white_flower_dish | white |
| white_half_ball | white |
| white_star | white |
| yellow_bin_top | yellow |
| yellow_bowl | yellow |
| yellow_ice_cream | yellow |
| yellow_plane | yellow |
| yellow_ship | yellow |
| yellow_star | yellow |
| yellow_tree | yellow |
| blue_foot | blue |
| blue_pot | blue |
| blue_shell | blue |
| blue_tower | blue |
| brown_flower_dish | brown |
| erdal_lid | mixed |
| golden_lid | golden |
| green_apple | green |
| green_half_ball | green |
| grey_manner_minder_front | grey |
| orange_lid | orange |
| orange_tower | orange |
| pink_crab | pink |
| pink_half_ball | pink |
| pink_seahorse | pink |
| red_cup | red |
| white_ashtray | white |
| white_box | white |

| | |
|---|---|
| yellow_egg_holder | yellow |
| yellow_shovel | yellow |
| yellow_strawberry | yellow |
| green_aloe_vera_box | green |
| black_coffee_filter | black |
| blue_berry | blue |
| blue_crab | blue |
| blue_cup | blue |
| brown_pen_holder | brown |
| golden_cup | golden |
| green_pot | green |
| green_banana | green |
| light_blue_train | blue |
| silver_coffee_pot | silver |
| pink_ship | pink |
| red_tube | red |
| silver_tape_(seminar_room) | silver |
| white_orbit_box | white |
| white_wheel | white |
| yellow_lid | yellow |
| yellow_lobster | yellow |
| light_blue_tower | blue |
| silver_coffee_box | silver |
| brown_flower_pot | brown |
