## Supplementary Table 1 (subjects) for "Going back to ‘basics’: Harlow’s learning set task with wolves and dogs"

| Name | Species | Sex | Experiments | Age at Exp.1 (months) | Last Exp.1 set | Age at Exp.2 (months) | Last Exp.2 reversal |
|---|---|---|---|---|---|---|---|
| **Alika** | dog | f | Exp.1 | 10 | 8 | | |
| **Asali** | dog | m | Exp.1 & Exp.2 | 13 | 45 | | |
| **Banzai** | dog | m | Exp.1 | 9 | 9 | | |
| **Bashira** | dog | f | Exp.1 | 13 | 16 | | |
| **Binti** | dog | f | Exp.1 & Exp.2 | 13 | 28 | 35 | 37 |
| **Bora** | dog | f | Exp.1 | 12 | 54 | | |
| **Enzi** | dog | m | Exp.1 | 9 | 9 | | |
| **Gombo** | dog | m | Exp.1 | 10 | 15 | | |
| **Hakima** | dog | m | Exp.1 & Exp.2 | 13 | 28 | 33 | 2 |
| **Hiari** | dog | m | Exp.1 | 10 | 17 | | |
| **Imara** | dog | f | Exp.1 | 10 | 10 | | |
| **Kilio** | dog | m | Exp.1 & Exp.2 | 9 | 31 | 22 | 29 |
| **Layla** | dog | f | Exp.1 | 12 | 43 | | |
| **Maisha** | dog | m | Exp.1 & Exp.2 | 9 | 25 | 21 | 72 |
| **Meru** | dog | m | Exp.1 & Exp.2 | 22 | 49 | 52 | 7 |
| **Nia** | dog | f | Exp.1 | 10 | 51 | | |
| **Nuru** | dog | m | Exp.1 & Exp.2 | 10 | 36 | 40 | 14 |
| **Panya** | dog | f | Exp.1 | 22 | 12 | | |
| **Pepeo** | dog | m | Exp.1 | 23 | 9 | | |
| **Rafiki** | dog | m | Exp.1 | 61 | 55 | | |
| **Sahibu** | dog | m | Exp.1 | 10 | 6 | | |
| **Zuri** | dog | f | Exp.1 | 10 | 60 | | |
| **Amarok** | wolf | m | Exp.1 & Exp.2 | 10 | 20 | 29 | 24 |
| **Apache** | wolf | m | Exp.1 & Exp.2 | 14 | 40 | 34 | 3 |
| **Aragorn** | wolf | m | Exp.1 & Exp.2 | 9 | 15 | 19 | 88 |
| **Cherokee** | wolf | m | Exp.1 | 10 | 47 | | |
| **Chitto** | wolf | m | Exp.1 | 10 | 24 | | |
| **Geronimo** | wolf | m | Exp.1 & Exp.2 | 10 | 63 | 43 | 32 |
| **Kaspar** | wolf | m | Exp.1 & Exp.2 | 9 | 16 | 13 | 144 |
| **Kay** | wolf | f | Exp.1 | 10 | 9 | | |
| **Kenai** | wolf | m | Exp.1 & Exp.2 | 11 | 41 | 38 | 40 |
| **Nanuk** | wolf | m | Exp.1 | 17 | 68 | | |
| **Shima** | wolf | f | Exp.1 & Exp.2 | 29 | 57 | 28 | 80 |
| **Tala** | wolf | f | Exp.1 | 29 | 45 | | |
| **Tatonga** | wolf | f | Exp.1 | 45 | 50 | | |
| **Una** | wolf | f | Exp.1 | 9 | 20 | | |
| **Wamblee*** | wolf | m | Exp.1 | 11 | 29 | | |
| **Wapi** | wolf | m | Exp.1 | 11 | 38 | | |
| **Yukon** | wolf | f | Exp.1 & Exp.2 | 15 | 20 | 23 | 46 |

*: Date for first session not recorded, date for second session used to calculate age.
